# Identification of a new Inovirus in the absence of bacterial infection in the human virome

**DOI:** 10.1101/2022.03.09.483013

**Authors:** Nikolay Popgeorgiev, Mart Krupovic, Julien Hiblot, Laura Fancello, Sonia Monteil-Bouchard, Christelle Desnues

## Abstract

Viruses infecting bacteria, known as bacteriophages, represent the most abundant viral particles in the human body. They participate in the control of the human associated bacterial communities and play an important role in the dissemination of virulence genes. Here we present the identification of a new genetic element, named RIP1, in the human virome. RIP1 shares conserved structural genes with the single-stranded DNA viruses of the *Inoviridae* family. Furthermore structure-function studies identified nuclear subcellular localization of the RIP1 DNA replication/recombination machinery. Additional metagenomics analysis and polymerase chain reaction detected RIP1 in multiple body sites including blood, cerebrospinal pericardial and amniotic fluids, nasal swabs and feces in the absence of concomitant bacterial infection, uncovering inovirus phage persistence in the human virome.

## Introduction

The human body is the home of diverse viral flora^1^. Viruses that infect bacteria, also known as bacteriophages, are the most abundant and diverse viral entities in the human body^2^. Bacteriophages are detected in virtually all anatomical sites, being most abundant in the digestive system^3^, the respiratory tract^4^ and on the skin^5^. Phage communities play a critical role in the control of the bacterial populations in humans. More recently bacteriophages have also emerged as potential virulence gene carriers, which can participate in the bacterial pathogenicity through lateral gene transfer. Seminal observations in viral metagenomes of the oral cavity of healthy individuals as well as cystic fibrosis patients showed that phages represent an important reservoir for bacterial virulence and resistance genes, thus contributing to bacterial pathogenicity ^4,6,7^.

Filamentous bacteriophages belong to the genus *Inovirus* (family *Inoviridae*) and infect mostly Gram-negative bacteria^8^. The phage virions are slender filaments usually about 6 nm in diameter and 800–2000 nm long. Inovirus genomes consist of circular single-stranded DNA (ssDNA) molecules of about 4.5–12.4 kb, which typically display a modular organization, with genes encoding proteins responsible for genome replication, virion morphogenesis, and structure compacted into clusters^9^. They replicate *via* a rolling-circle (RC) mechanism initiated by the phage-encoded replication initiation protein (REP). In humans, inoviruses are known to contribute to the virulence of pathogenic *Vibrio cholerae* strains^10^. Indeed, chromosomally integrated CTX□ prophage encodes a suite of toxins, including the primary cholera toxin (genes *ctxA* and *ctxB*), which causes watery diarrhea. Two additional proteins with enterotoxic activity—the zonular occludens toxin (ZOT) and accessory cholera enterotoxin (ACE)—increase short-circuit current across rabbit intestinal tissue by altering tight junctions, allowing the passage of macromolecules through mucosal barriers and thereby contributing to *Vibrio cholera* pathogenicity^10^. Whereas CtxAB toxin is not essential for CTX□ reproduction, ZOT and ACE are required for virion assembly and structure, respectively. Notably, due to its critical role in phage assembly, a homologue of ZOT is also conserved in inoviruses infecting a variety of hosts other than *Vibrio cholera*.

In the present study we present the identification and characterization of a new inovirus, named RIP1. RIP1 genome was detected using high-throughput DNA sequencing of samples obtained from various anatomical sites in the absence of concomitant bacterial infection or contamination, suggesting possible phage persistence in the human virome.

## Results

### Identification of a new Inovirus in the human virome

We have recently performed a comprehensive study of the viral composition in the human blood using a viral metagenomics approach^11^. During the course of this study we detected the presence of metagenomic reads homologous to the ssDNA *Ralstonia solanacearum* phages RSM1 and RSM3 (later referred to as RSM 1/3), which are members of the *Inoviridae* family. *De novo* genomic assembly of the blood metagenome revealed that these reads organized a single contig of 8,5 kb in length with 16 predicted open reading frames (ORF) (Fig.1A). We identified several *cis*-regulatory elements typical of inoviruses present in the RIP1 genome. Among these we detected a conserved *attP* core sequence (between 1846-1858) required for DNA integration/recombination and which was previously identified and functionally characterized in RSM 1/3 phages^12^ (Fig.1A). At the 3’ end of the contig we identified a long hairpin that potentially corresponds to the phage packaging signal typically found in filamentous phages (Fig.1A). Notably, we also detected a tandem repeat sequence containing only 1 internal mismatch at position 2740-2974 with unknown function.

**Figure 1:**
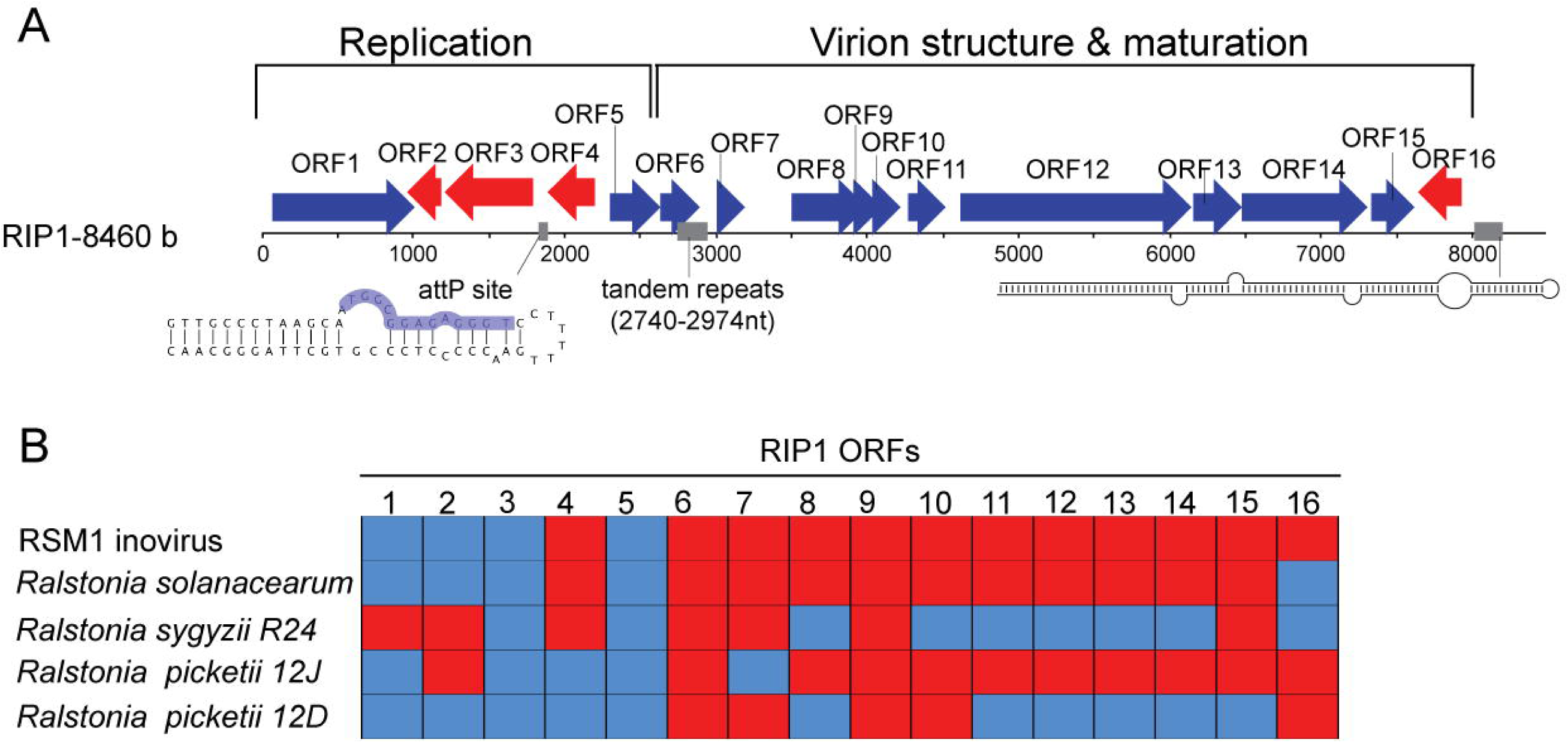
Identification of a new Inovirus RIP1. **(A)** Linear representation of the genomic organization of RIP1 element. The length in bases (b) was given on the left. Forward/reverse ORFs are represented with blue/red arrows respectively. The presence of *cis* elements are outlined with grey boxes. The secondary structure of the *attP* site and the long hairpin, corresponding to the predicted genome packaging signal, are represented below the linearized sequence. The core sequence of the *attP* site is outlined in blue. **(B)** The presence/absence of homologous genes in *Ralstonia picketii 12D, Ralstonia sygyzii R24, Ralstonia picketii 12J, Ralstonia solanacearum CRM15* and RSM1 phage genomes is reported using blue/red boxes, respectively.

The majority of RIP1 ORFs coded for hypothetical proteins and had their homologous counterparts in the RSM 1/3 viral genomes as well as in the inovirus-like prophage genomes integrated in bacterial chromosomes of the *Ralstonia* group (Fig.1B&S1). Thus, we named this metagenomic contig RIP1 (Ralstonia Inovirus phage 1). BlastP analyses and thorough comparison of RIP1 ORF products with those encoded by other phages allowed identification of all structural proteins typical of inoviruses among the gene products of RIP1 (Table 1). Indeed, ORF11 encodes for pVIII-like (here and elsewhere the protein nomenclature of M13-like inoviruses is used) viral major capsid protein, which organizes into a helical array covering the phage DNA and forming the virion. RIP1 ORF11 protein contains the characteristic N-terminal signal sequence, followed by amphiphatic, hydrophobic and basic domains (Table1; Fig.S2A), essential for positioning of the protein in the membrane prior to virion assembly, tight packing of the capsid proteins in the virion tube following the assembly and interaction with the phage DNA within mature virions^8^. RIP1 ORF9 and ORF10 were respectively identified as the homologs of pVII and pIX minor capsid proteins of M13-like inoviruses. The two proteins are located at the tip of the filamentous phage particle, which is the first to emerge during virus assembly; both are small, roughly 3-3.5 kDa, membrane proteins and possess 1 transmembrane domain each (Table1; Fig.S2B&C). ORF12 and ORF13 are homologous to the pIII-like and pVI-like (Fig.S2D) inoviral proteins, which mediate virion assembly termination, release and infection. pIII homologue in RIP1 was identified after several PSI-BLAST iterations using ORF12 sequence as a query. ORF5 corresponds to pV, which binds to single stranded form of the genome, thereby controlling the switch from double-stranded replicative intermediate to the synthesis of the (+) stand, which is subsequently packed into the progeny virions. RIP1 ORF14 corresponds to pI, which is homologous to the ZOT toxin encoded by CTX□ phage. However, the C-terminal region of the CTX□ ZOT protein responsible for its activity as a toxin (288–293 aa) was lacking in the RIP1 ORF14.

**Table1:** Summary information about the predicted ORFs of RIP1 phage. From the left to the right, ORF number, location, orientation, corresponding protein length, conserved domains and predicted functions were reported. The research of conserved domains and signatures were performed using CDD, PFAM and Prosite databases.

| ORF# | Position (bases) | Strand | Protein length | Conserved domains | Predicted functions |
| --- | --- | --- | --- | --- | --- |
| 1 | 62-1012 | + | 316 | NOT FOUND | rolling circle DNA replication initiation (pII-like) |
| 2 | 963-1187 | - | 74 | NOT FOUND | Unknown (Hypothetical protein) |
| 3 | 1203 - 1793 | - | 196 | Resolvase, N terminal domain (2-140aa); Helix-turn-helix domain (148 -184aa) | DNA recombination (Resolvase-like) |
| 4 | 1895 - 2197 | - | 100 | Helix-turn-helix domain (16-61 aa); DUF4072 domain (18-47 aa) | DNA binding |
| 5 | 2302 - 2631 | + | 109 | Helix-destabilising domain (10-35aa); Phage DNA bind | ssDNA binding (pV-like) |
| 6 | 2628 - 2894 | + | 88 | NOT FOUND | Protein folding, Chaperone (Highly divergent DNA J-like protein) |
| 7 | 3002 - 3190 | + | 62 | AA transfer class 2 attachment site (20-29aa) | Unknown (Hypothetical protein) |
| 8 | 3495 - 4001 | + | 168 | GrpE family [PF01025] (21-104aa) | Protein folding, Chaperone (GrpE-like) |
| 9 | 4012 - 4098 | + | 28 | Hydrophobic domain (5-27) | Phage assembly, minor capsid protein (pVII-like) |
| 10 | 4095 - 4214 | + | 39 | Transmembrane domain (11-32aa) | Phage assembly, minor capsid protein (pIX-like) |
| 11 | 4272 - 4523 | + | 83 | N-terminal signal sequence (30), amphiphatic (40-54), hydrophobic (55-71) and basic domains (72-83) | Phage assembly (pVIII-like) |
| 12 | 4617 - 6137 | + | 506 | Threonine-rich region (303 – 364aa) | Termination of virion assembly (pIII-like) |
| 13 | 6150 - 6479 | + | 109 | Hydrophobic domains (4-20 ;33-55 ;75-97) | Termination of virion assembly (pVI-like) |
| 14 | 6481 - 7311 | + | 276 | Zonular occludens toxin (ZOT) domain | Phage assembly (pI-like) |
| 15 | 7334 - 7618 | + | 94 | NOT FOUND | Unknown (Hypothetical protein) |
| 16 | 7640 - 7927 | - | 95 | NOT FOUND | Unknown (Hypothetical protein) |

### RIP1 ORF1 and ORF3 possess nuclear subcellular localization

We next focused on the RIP1 ORF1 and ORF3. ORF1 encodes for a REP homolog required for phage DNA replication through rolling-circle (RC) mechanism. All three signature motifs found in RC-REP proteins of bacterial, archaeal and eukaryotic ssDNA viruses^13,14^ were conserved in the product of ORF1 (Fig. S3). ORF3 encodes a resolvase-like serine recombinase (RES) required for phage integration in the bacterial genome (Fig. S4). Notably, the integration/excision activity of the close homologue of RIP1 ORF3 from RSM1 phage has been demonstrated experimentally^15^. Unexpectedly, using NLStradamus web based interface, we detected a eukaryotic nuclear localization signal (NLS) in the REP protein (Fig.2A). To confirm this observation, we expressed REP protein in human embryonic kidney (HEK) cells. Immunofluorescence experiments indeed detected REP protein in the nucleus (Fig.2B,C). Furthermore, RES protein also showed similar subcellular localization although NLS was not detected in its primary structure (Fig.2C). Of note, nuclear localization was not systematic for all RIP1 proteins. Indeed, expression of the ZOT protein (ORF14), required for the phage maturation, in HEK cells resulted in cytoplasmic localization, suggesting that the nuclear targeting is restricted to the phage DNA replication/recombination machinery.

**Figure 2:**
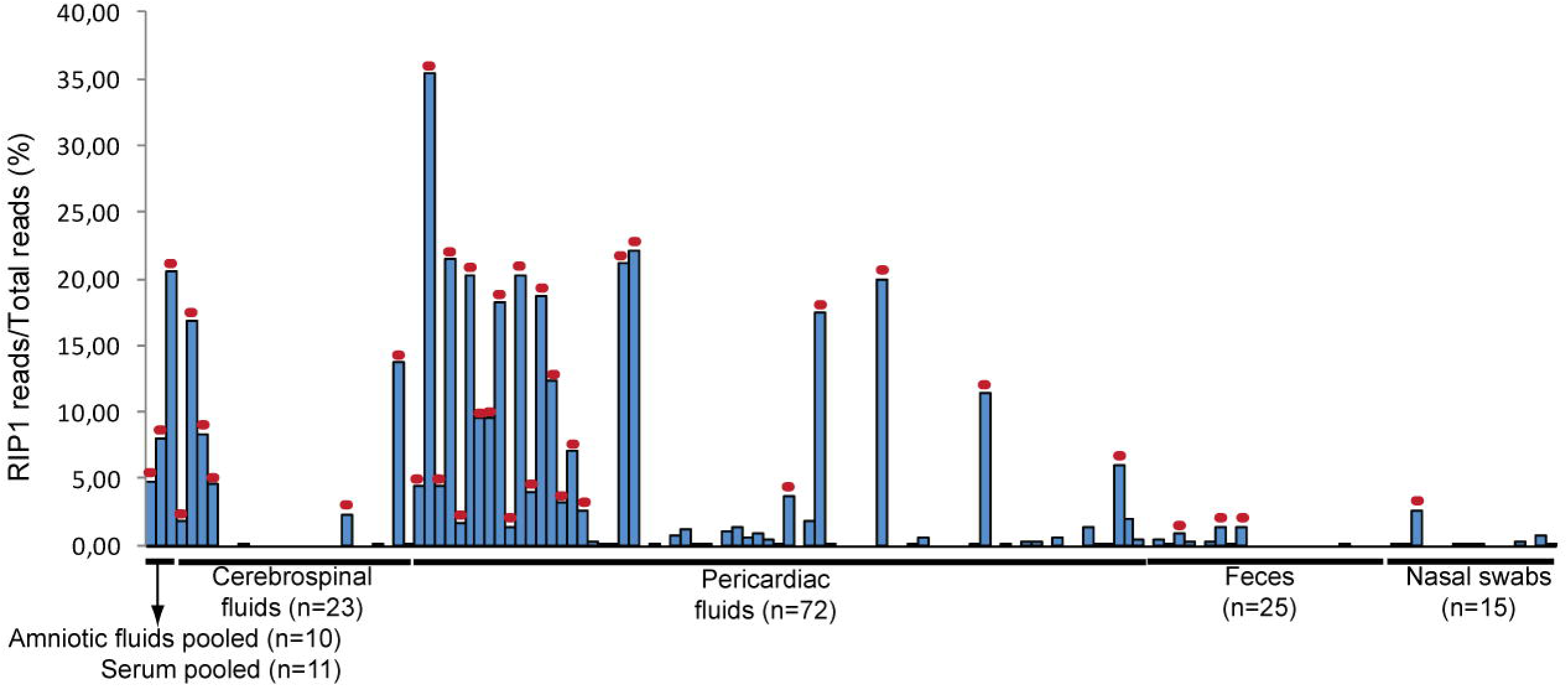
RIP1 REP and RES proteins have nuclear localization in human cells. (**A**) Primary and secondary structure of the REP protein. The predicted nuclear localization signal is highlighted with red boxes. Alpha helices are represented with blue cylinders, beta sheets are represented with orange arrows and unstructured regions with orange asterisks. **(B)** Western blot using anti-Flag antibody showing the expression of Flag-tagged REP, RES and ZOT proteins. Anti-ATPase antibody was used for calibration purposes. **(C)** Confocal images showing the subcellular localization of Flag-tagged REP, RES and ZOT proteins. Nuclei were stained using DAPI. Scale bars; 2µm. **(D)** Histogram presenting the percentage of cells having anti-Flag fluorescence signal co-localisation with DAPI nuclear dye (mean±s.d.; three independent experiments).

### RIP1 prevalence in the human virome is not concomitant with a bacterial infection

We further performed BlastN/X searches against nt/nr Genbank databases and genomic mapping on human metagenomes (n=138) generated in our lab from diverse anatomical sites, including amniotic, pericardiac and cerebrospinal fluids, feces and nasal swabs. We found that 77 (55.8%) of these metagenomes contain sequences matching to RIP1 genome. The number of RIP1-specific reads was more significant in the human fluid samples, representing up to 35% of the total metagenomics reads (Fig.3). Importantly, additional tBlastN searches against expressed sequence tags (EST) and transcriptome shotgun assembly (TSA) databases using RIP1 ORFs as queries identified transcripts corresponding to the ORF1, ORF2 and ORF3 in other animals, including *Sus scofa, Schistosoma mansoni* and *Tupaia chinensis*, suggesting that the presence of RIP1-related viruses is not restricted to humans (Table S1).

**Figure 3:**
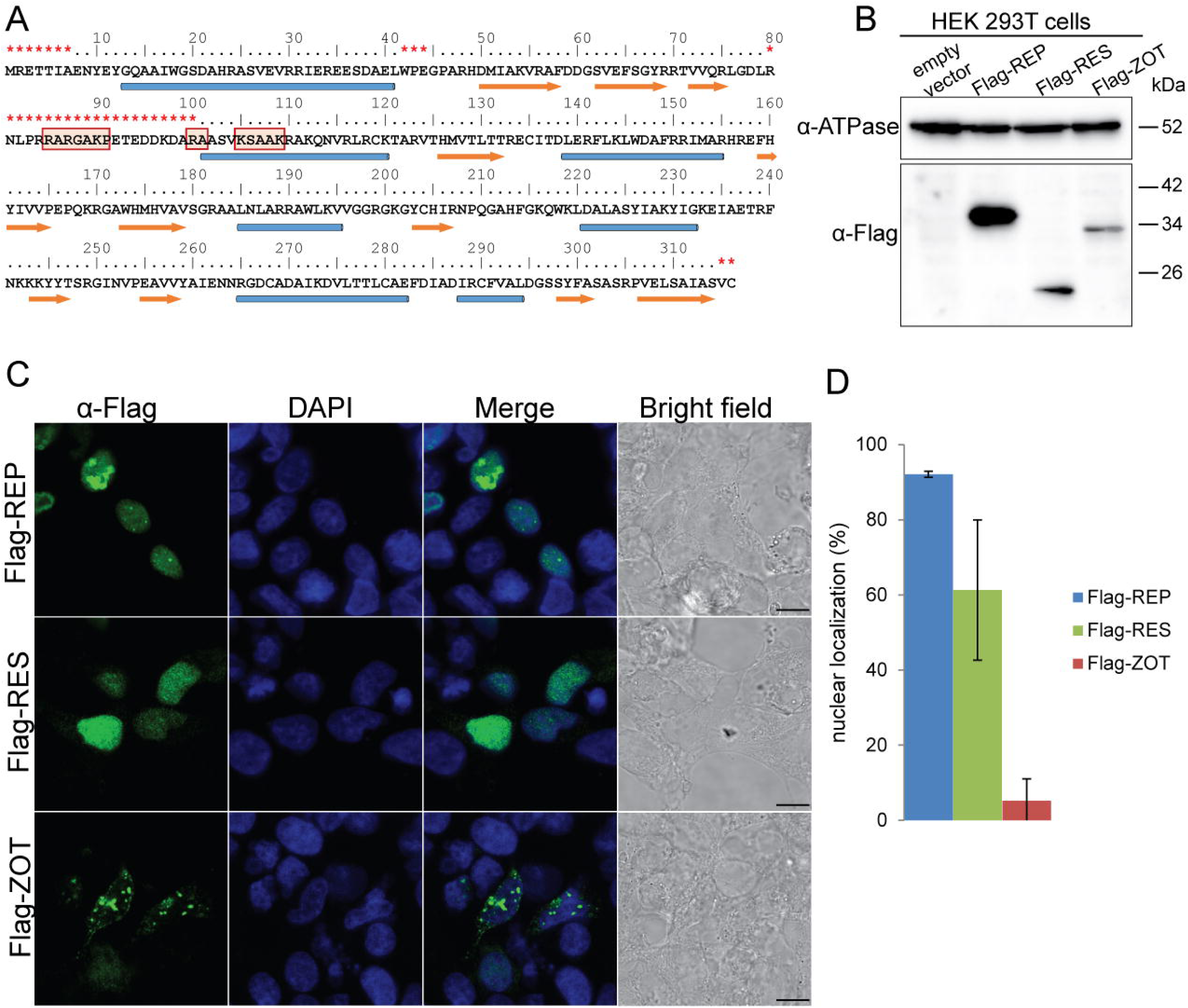
RIP presence in the human virome. Histogram showing the fraction of metagenomics reads corresponding to RIP genome found in different human samples. Samples from which *de novo* prediction was possible are outlined with red dots.

Recently, the discovery of a ssDNA virus genome, which was initially assumed to be associated with human samples, but subsequently traced to a contamination of the DNA extraction spin columns was reported^16^. The formal possibility that the presence of RIP1 genome in human samples was due to contamination was excluded by systematic testing of all kits and reagents used in this study for DNA isolation and amplification by polymerase chain reaction (PCR). This analysis showed that none of the reagents and kits contained RIP1 DNA. Furthermore, the presence of RIP1 in the viral metagenomes was verified by performing PCR using ORF14 specific primers on viral DNA extracted from sera samples from blood donors. Notably, metagenomics analysis as well as pan-genomic 16S rRNA and *Ralstonia pickettii* specific 16S rRNA PCRs did not reveal bacterial presence in these samples, suggesting that RIP1 detection was not concomitant with a bacterial infection.

## Discussion

Here we present the identification of a new bacteriophage named RIP1 in the human virome. Genome organization as well as genetic composition strongly suggests that RIP1 is related to bacteriophages of the *Inoviridae* family. All components required for the assembly and structure of M13-like inoviruses are conserved in the RIP1 genome (Fig. S2). Based on the comparison of the RIP1 related sequences, it appears that it shares the most recent common ancestor with proviruses integrated in the genomes of *Ralstonia pickettii* 12D and *Ralstonia syzygii* R24 (Fig. 1B). Notably, in all three elements, the ZOT-like proteins lack the toxigenic peptide found in the C-terminal region of the ZOT protein of the *Vibrio cholera* phage CTX□^17^, suggesting that the proteins of *Ralstonia* phages are unlikely to elicit adverse effects on the tight junctions.

Comparative genomic analysis of the RIP1 genome revealed that it is a mosaic of genes derived from various inoviruses (Fig. S1). Indeed, genomes of the *Inoviridae* members are known to be shaped by frequent recombination events, which often lead to non-orthologous gene replacements and acquisition of new genes^9^. Thus, the RIP1 ancestor has apparently emerged as a result of recombination between an RSM1-like phage, which has donated the genome replication/recombination modules, and a phage that contributed the virion structure and assembly module.

Strikingly, we found that 55.7% of the analysed metagenomes contain RIP1 sequences, which in some cases represented up to one third of all metagenomics reads. However, viral metagenomes were amplified using phi29 DNA polymerase, which displays preference towards circular ssDNA matrices^18^, thus rendering RIP1 abundance estimation difficult. RIP1 occurrence in multiple human samples points towards temporal phage persistence after asymptomatic infection. Indeed, the most likely route for RIP1 emergence in the human virome is *via* its *Ralstonia* host. Human-associated *Ralstonia pickettii* were found to be part of the commensal flora of the oral cavity and the upper respiratory tract of healthy individuals^19,20^. Furthermore, *Ralstonia pickettii* can be isolated from a variety of clinical specimens, including sputum, blood, wound infections, urine, ear and nose swabs, and cerebrospinal fluids^21^. In this study 16S rRNA gene PCRs were unable to identify the presence of bacterial infection in our samples, thus strongly suggesting that RIP1 detection is not due to bacterial contamination. Another possibility, even though highly speculative, is that REP- and/or RES-mediated introduction of the RIP1 genome into the nucleus could lead to occasional proliferation of RIP1 in the human genome in the form of a mobile genetic element.

Bacteriophages are generally not considered to have any direct effect on eukaryotes. Recently, however, terminal proteins, which are covalently attached to the termini of the genomes of certain dsDNA bacteriophages, were shown to contain functional NLS^22,23^. This finding has suggested that the genomes of these bacteriophages have an access to the eukaryotic genomes and might participate in shaping of the latter by horizontal gene transfer. In the present study we have shown that proteins of an ssDNA inovirus that are involved in DNA metabolism, including replication initiation (RC-REP) and recombination (RES), also can enter the eukaryotic nucleus. Importantly, upon replication initiation RC-REPs form a covalent intermediate with the viral DNA, suggesting that like in the case of terminal protein-encoding bacteriophages^22^ RIP1 REP might shuttle the inoviral DNA into the nucleus. This opens an intriguing possibility that RIP1 proteins might elicit their corresponding enzymatic activities in the context of eukaryotic genomes. For example, RES could mediate recombination between the phage and cellular chromosomes. Notably, serine recombinases are widely used in genetic engineering for stable integration of transgenes into eukaryotic cells^24^. Interestingly, another RIP1 protein with functional NLS is homologous to the RC-REPs of eukaryotic ssDNA viruses and prokaryotic TnpA-like transposases of the IS200/IS605 family^25^ (Fig. S3). Furthermore, certain classes of eukaryotic transposons are believed to have recently evolved from parvoviruses and geminiviruses^26^, suggesting that the two classes of genetic parasites, transposons and viruses, occasionally interconvert. It is thus tempting to speculate that prokaryotic ssDNA viruses, such as RIP1, might give (or even have given) rise to new classes of eukaryotic transposable elements. Additionally, recombination between different groups of ssDNA viruses has played a key role in the evolution of this virus class^14^. Co-localization of prokaryotic and eukaryotic ssDNA viruses thus provides additional opportunities for exploring the genetic landscape for generating novel recombinant virus types. Obviously, further studies will be required to clarify the origin, function and impact of RIP1 in the human virome.

## Methods

### Human samples

A total of 138 samples collected from blood serum, amniotic, pericardiac, cerebrospinal fluids, feces and nasal swabs were used in this study. All samples were collected between 2007 and 2010 in 5 French hospitals (the Hospital of Niort and La Timone, Nord, Conception and Clairval hospitals in Marseille). Sera samples (n=11) were collected at the “Etablissement Français du Sang” Marseille. Sera and amniotic fluids were pooled prior DNA extraction.

### Ethics Statement

All samples used in this study were collected from human subjects using a protocol approved by the local ethics committee IFR48 (Marseille, France). Written informed consent was obtained from the parents or legal guardians of all subjects.

### Viral isolation and high-throughput sequencing

Each sample was centrifuged at low speed to eliminate proteins and cellular debris. The resulting supernatant was collected and filtered through 0.45-µm filter pore. Virus-like particles were concentrated by ultracentrifugation at 55,000 g for 60 min. The resulting pellet was resuspended in a phosphate buffered saline solution (PBS) previously filtered at 0.02 µm. Purified VLPs were treated with DNase and RNase to remove any residual host and bacterial DNA as previously described^27^. Viral DNA was then extracted using the High Pure Viral Nucleic Acid Kit (Roche Applied Science) following the manufacturer’s recommendations. Extracted DNA was amplified using the commercial Illustra™ GenomiPhi V2 DNA Amplification Kit (GE Healthcare Life Sciences) to generate sufficient material for shotgun 454 pyrosequencing library preparation. Amplified DNA was purified using Agencourt AMPure XP-PCR Purification kit (Beckman Coulter) to remove the enzyme, dNTPs and primers, and subsequently sequenced on a 454 Life Sciences Genome Sequencer FLX instrument using titanium chemistry (Roche Applied Science).

### Reads processing, mapping, *de novo* assembly and ORF prediction

Obtained sequences from 454 pyrosequencing were screened to remove exact and nearly identical duplicates. Duplicate removal was performed by the CD-HIT-454 program available under the CAMERA 2.0 web portal. Mapping of metagenomic reads and *de novo* assembly were performed using the CLC Genomics Workbench version 4.9 (www.clcbio.com). Mapping onto *Ralstonia pickettii* 12D chromosome 1 reference genome (Acc. Number # NC_012856.1) was performed with a minimal overlap length fraction of 0.5 and a minimal similarity of 0.95 as mapping parameters. *De novo* contig assemblies were performed with a minimum overlap length fraction of 0.5 and a minimum overlap identity of 0.9. Open Reading Frames prediction was performed using Prodigal software.

### Polymerase chain reaction and sequencing

Standard PCR amplification was performed using Phusion High-Fidelity DNA Polymerase (Thermo Scientific). The absence of kit/reagent contamination was verified in the High Pure Viral Nucleic Acid Kit (Roche Applied Science) and Illustra™ GenomiPhi V2 DNA Amplification Kit (GE Healthcare Life Sciences) using 0.02 µm filtered PBS buffer as a sample. PCR amplifications were performed using RP1F/ RP1R and RP3F/ RP3R primers with the following parameters: an initial denaturation step at 98 °C for 30 s, followed by 35 cycles of 98 °C for 30 s, 60 °C for 30 s, and 72 °C for 45 s. The presence of *Ralstonia pickettii* 16S rRNA gene was verified using Rp-F1/Rp-R1 forward/reverse primers as previously described^19^. To confirm the presence of sequences found *in silico*, standard PCRs targeting RIP1 ZOT were performed using ZotIntF1/ZotIntR2 primers with the following parameters: an initial denaturation step at 98 °C for 30 s, followed by 35 cycles of 98 °C for 30 s, 52 °C for 30 s, and 72 °C for 20 s.. PCR positive samples were sequenced using the BigDye Terminator v1.1 Cycle Sequencing Kit (Life Technologies) according to the manufacturer’s instructions. Primer sequences are presented in the supplementary table S1.

### RIP1 CDS cloning

ORF1, ORF3 and ORF14 ORFs were cloned in frame with the flag-tag at the 5’ region in pCS2+ expression vector using FlagRepF/FlagRepR, FlagResF/FlagResR, FlagZotF/FlagZotR primers respectively. For this purpose PCR amplicons and pCS2+ vector were digested using XhoI/BamHI restriction enzymes. Ligation was performed using T4 ligase (Promega) following manufacturer recommendations.

### Cell culture, transfection, immunofluorescence and western blot

Human embryonic kidney (HEK293T) cells were maintained under standard culture conditions (37°C and 5% CO2) and cultured in MEM media supplemented with 10% FBS, 1% Glutamine and 1% PS. Cells were transfected using Lipofectamine 2000 (Life Technologies) following manufacturer specifications. For cell immunofluorescence staining transfected cells were fixed 20 min in PFA 4% at 37°C then washed three times in PBS and incubated 20 min with blocking buffer (PBS + 0.1% Triton X-100, BSA 3%). Mouse FlagM2 antibody (Sigma-Aldrich) diluted at 1/200 in blocking buffer were incubated for 1H at room temperature followed by three washes in PBS + 0.1% Triton X-100. Secondary anti-mouse Alexa 488 antiboby (1/1000) was incubated for 1 hour. Cell nuclei were stained with ProLong Gold Antifade reagent (Molecular Probes) containing DAPI. For wertern blot, cell proteins were extracted in RIPA buffer (1% NP-40, 0.5% deoxycholic acid, 0.1% SDS in PBS, containing protease inhibitors). Proteins were the resolved in a 12% SDS-PAGE gel and transferred on a 0.2 µm nitrocellulose membrane and then immunobloted using mouse FlagM2 antibody (Sigma-Aldrich) diluted at 1/500.

## Acknowledgments

This work was supported by the Starting Grant #242729 from the European Research Council to C. Desnues.

## Author contributions

N.P. and C.D. designed the experiments. N.P. and S.M. performed the experiments. N.P., M.K, L.F., D.R., J.H., and C.D. analysed the results. N.P. and M.K. wrote the paper.

## Figure Legends

**Figure S1:**
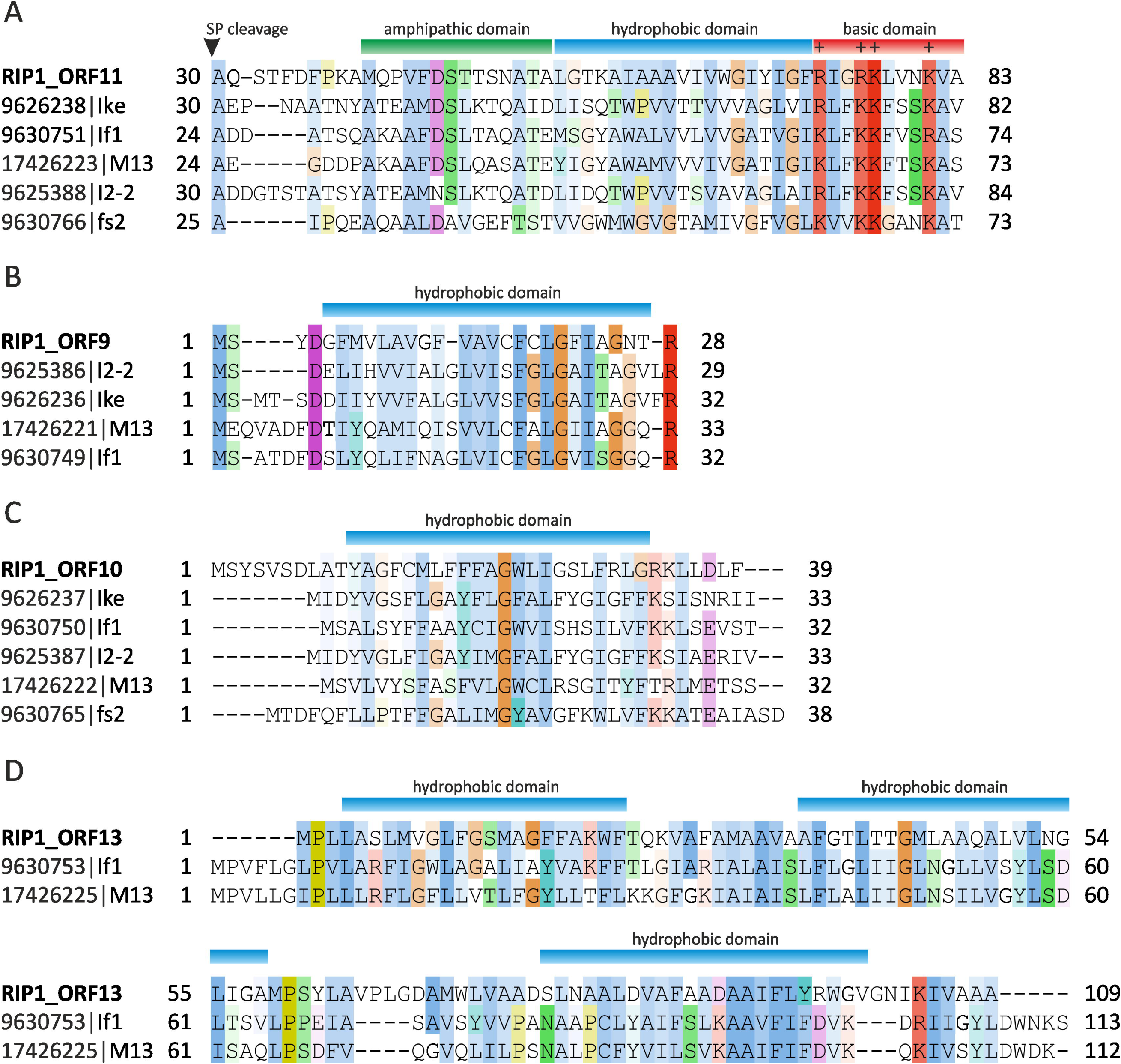
RIP1 has a chimeric genomic organization. (A) Schematic representation of local alignment results between RIP1 genome and genomes of *Ralstonia picketii 12D, Ralstonia sygyzii R24, Ralstonia picketii 12J, Ralstonia solanacearum CRM15* and RSM1 phage. The matching regions as well as their locations on the RIP1 genome are reported using black dashes. (B) Genomic organization comparison between RIP1 genome and homologous bacterial regions. Forward/reverse ORFs are represented with blue/red arrows respectively. The presence of *attP* sites are represented with yellow boxes. Best Blast hits correspondence between RIP1 ORFs and bacterial ORFs are represented with colored lines. The presence of *cis* elements are outlined with beige boxes. DNA sequence homologous to *Ralstonia sygyzii R24* non coding region located inside ORF6 is reported with orange boxes.

**Figure S2:**
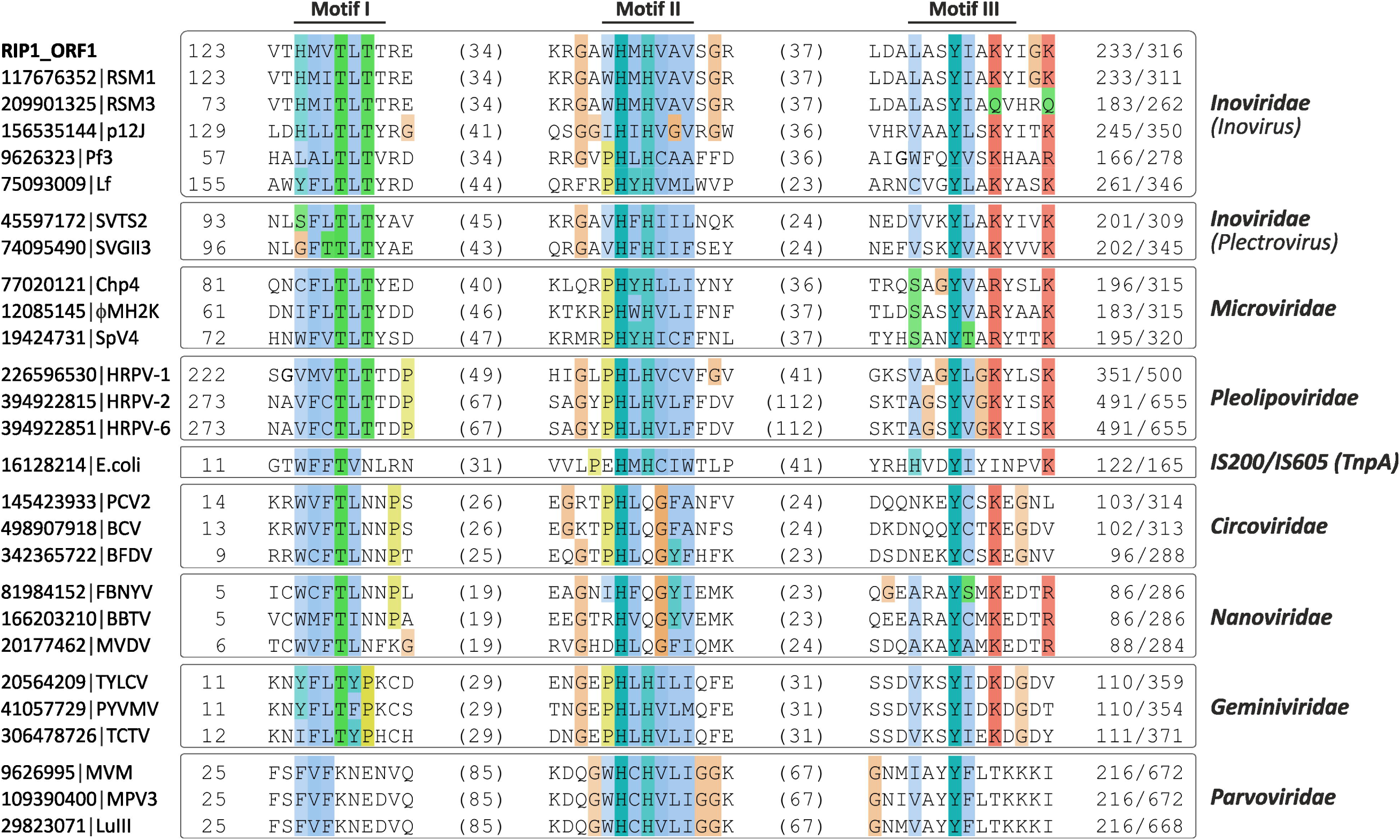
Multiple alignments between RIP1 ORFs and homologous inovirus proteins. **(A)** Multiple sequence alignment between ORF9 and pVII homologs of inoviruses. The location of a single hydrophobic domain is represented with a blue bar. **(B)** Multiple sequence alignment between ORF10 and pIX-like proteins of inoviruses. **(B)** Multiple sequence alignment between ORF11 and pVIII-like inoviral homologs. The amphiphatic, hydrophobic and basic domains are represented with green, blue and red bars, respectively. Positively charged residues are indicated with (+). (**D**) Multiple sequence alignment between ORF13 and pVI-like inoviral homologs. Abbreviations: SP, signal peptidase; Ike- Enterobacteria phage Ike; If1- Enterobacteria phage If1; I2-2- Enterobacteria phage I2-2; M13- Enterobacteria phage M13; fs2- Vibrio phage fs2. GenBank identifiers for the depicted proteins are provided in the figure.

**Figure S3:**
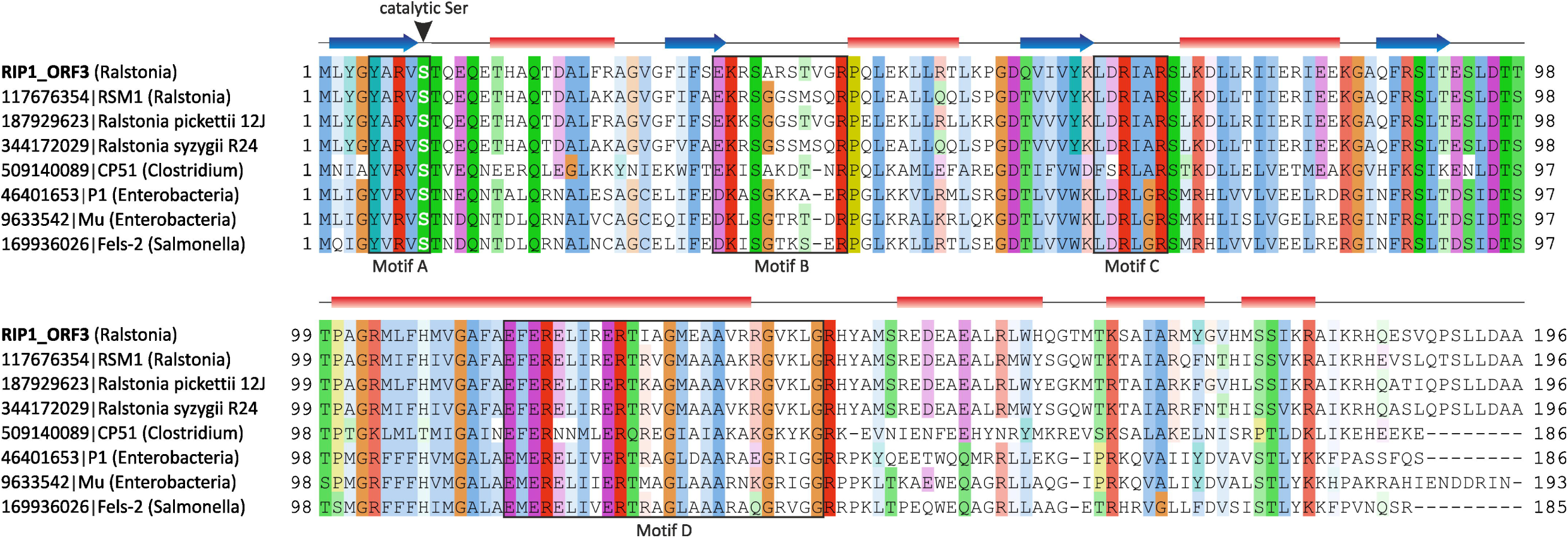
RIP1 ORF1 encodes a rolling-circle replication intiation protein. Alignment of the RC-REP protein encoded by RIP1 with the corresponding proteins encoded by ssDNA viruses infecting bacteria (*Inoviridae* and *Microviridae*), archaea (*Pleolipoviridae*), and eukaryotes (*Circoviridae, Nanoviridae, Geminiviridae and Parvoviridae*) as well as TnpA transposase of the IS200/IS605 family from *E. coli*. The protein sequences are denoted by their GenBank identiers, followed by the corresponding virus names. Only the three conserved motifs (I–III) typical of the RC-REPs are shown. The limits of the depicted motifs are indicated by the residue positions on each side of the alignment, while the number following the slash sign corresponds to the total length of the protein. The numbers in the parenthesis within the alignment indicate the distance between the motifs.

**Figure S4:**
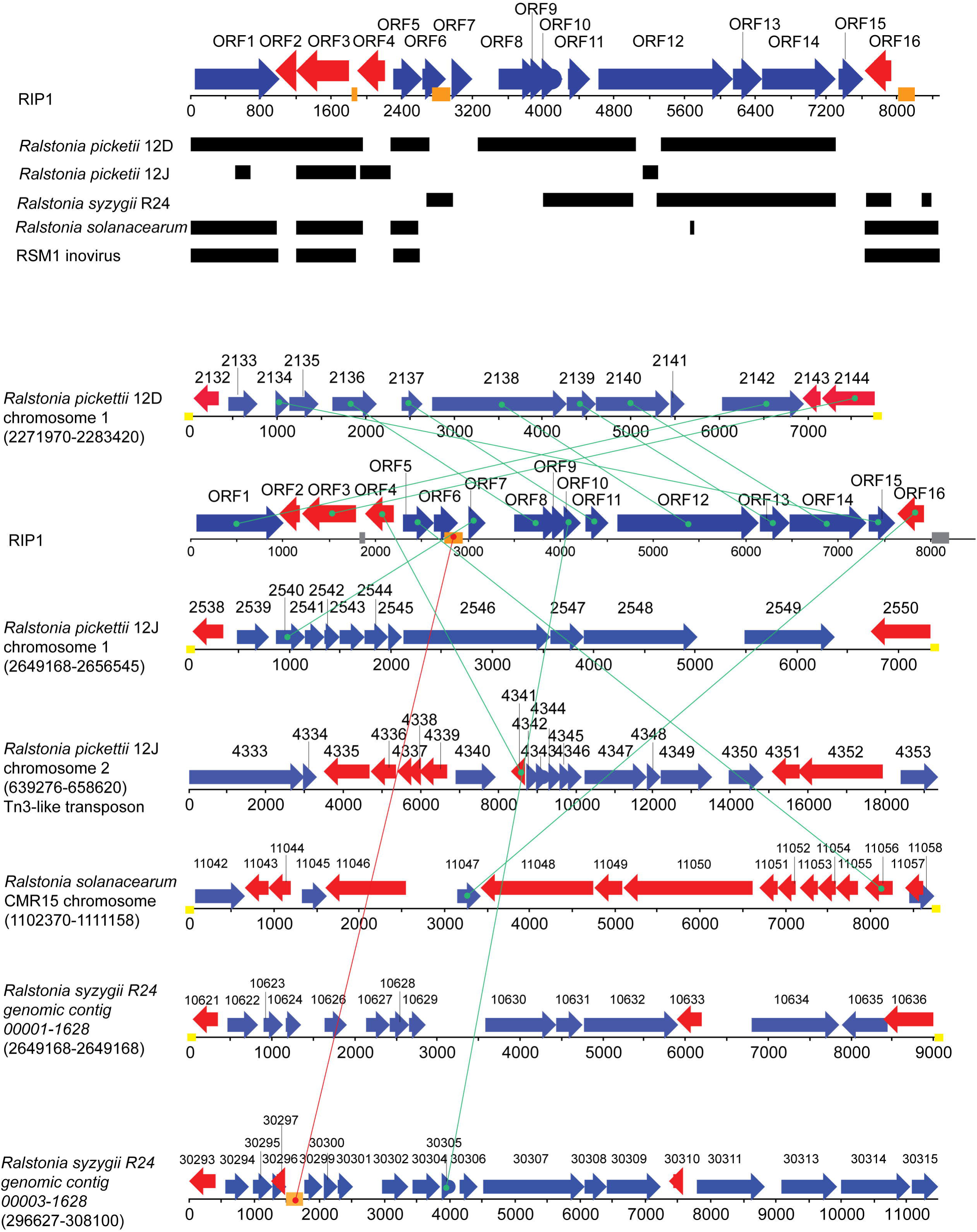
RIP1 ORF3 encodes a serine recombinase. Alignment of the RIP1 ORF3 product sequence with the corresponding sequences of serine recombinases encoded by various prophages and bacteriophages. The four conserved motifs (A–D) typical of serine recombinases are boxed, while the position of the catalytic serine residue is indicated with an arrowhead. The predicted secondary structure elements (α-helixes, red rectangles; β-strands, blue arrows) are shown above the alignment. Proviruses are indicated by the names of the organisms in which they reside, while the host names of the bacteriophages are provided in parenthesis. GenBank identifiers for all depicted proteins are provided in the figure.

