## Supplemental Table 1&2 for "Identification of a new Inovirus in the absence of bacterial infection in the human virome"

| **tBLASTn** | | | | |
| --- | --- | --- | --- | --- |
| RIP1 ORFs | Organism name | # Genbank | % Identity | E-value |
| ORF1 (REP) | *Sus scrofa* | gb\|CN025072.1\| | 95 | 8e^-19^ |
|  | *Schistosoma mansoni* | gb\|CF497648.1\| | 73 | 1e^-13^ |
| ORF2 | *Schistosoma mansoni* | gb\|CF497648.1\| | 59 | 3e^-13^ |
| ORF3 (RES) | *Schistosoma mansoni* | gb\|CF497648.1\| | 91 | 5e^-48^ |
|  | *Tupaia chinensis* | gb\|JU128735.1\| | 47 | 2e^-43^ |

**TableS1: tBlastN searches against EST and TSA databases using RIP1 ORFs**

| **Primer** | **Sequence (5’-3’)** |
| --- | --- |
| **FlagRepF** | ATATGGATCCATGGACTACAAGGACGACGATGACAAGATGCGTGAAACTACGATAGCAG |
| **FlagRepR** | ATATCTCGAGTCAACAAACACTGGCTATGGC |
| **FlagResF** | ATATGGATCCATGGACTACAAGGACGACGATGACAAGATGTTGTACGGTTACGCCCG |
| **FlagResR** | ATATCTCGAGTTACGCGGCGTCTAGCAAGC |
| **FlagZotF** | ATATGGATCCATGGACTACAAGGACGACGATGACAAGATGCCGATCTACATCATCACGG |
| **FlagZotR** | ATATCTCGAGTCACCATGAGGGGCATCGAG |
| **Rp-F1** | ATGATCTAGCTTGCTAGATTGAT |
| **Rp-R1** | ACTGATCGTCGCCTTGGTG |
| **ZotIntF1** | TGTGGATACGTTCACGCTCG |
| **ZotIntR2** | CGTTCGTCCAGTTTCGCTTC |
| **RP1F** | CCCGCAAGCGCGCCAACGCT |
| **RP1R** | ACGATCGGGATTGGTATCGC |
| **RP3F** | AAACAGTGCGTCGGTTTGTG |
| **RP3R** | ATCACGCGAGACGAACAACT |

**TableS2: Summary table of primers used in this study**
